# Deep learning-based molecular morphometrics for kidney biopsies

**DOI:** 10.1101/2020.08.23.263392

**Authors:** Marina Zimmermann, Martin Klaus, Milagros N. Wong, Ann-Katrin Thebille, Lukas Gernhold, Christoph Kuppe, Maurice Halder, Jennifer Kranz, Nicola Wanner, Fabian Braun, Sonia Wulf, Thorsten Wiech, Ulf Panzer, Christian F. Krebs, Elion Hoxha, Rafael Kramann, Tobias B. Huber, Stefan Bonn, Victor G. Puelles

**Author notes:** Equal contributions in first authors (Marina Zimmermann, Martin Klaus). Equal contributions in senior authors (Tobias B. Huber, Stefan Bonn, Victor G. Puelles). **Corresponding authors, Victor G. Puelles**, Postal address: III. Department of Medicine, University Medical Center Hamburg-Eppendorf; Martinistraße 52, Gebäude N27, 3. Etage, 20246 Hamburg, Germany, **Stefan Bonn**, Postal address: Center for Molecular Neurobiology Hamburg (ZMNH); Institute of Medical Systems Biology; Falkenried 94, 1. Etage, 20251 Hamburg, Germany, **Tobias B. Huber**, Postal address: III. Department of Medicine, University Medical Center Hamburg-Eppendorf; Martinistraße 52, Gebäude O10, 1. Etage, 20246 Hamburg, Germany.

## Abstract

Morphologic examination of tissue biopsies is essential for histopathological diagnosis. However, accurate and scalable cellular quantification in human samples remains challenging. Here, we present a deep learning-based approach for antigen-specific cellular morphometrics in human kidney biopsies, which combines indirect immunofluorescence imaging with U-Net-based architectures for image-to-image translation and dual segmentation tasks, achieving human-level accuracy. In the kidney, podocyte loss represents a hallmark of glomerular injury and can be estimated in diagnostic biopsies. Thus, we profiled over 27,000 podocytes from 110 human samples, including patients with anti-neutrophil cytoplasmic antibody-associated glomerulonephritis (ANCA-GN), an immune-mediated disease with aggressive glomerular damage and irreversible loss of kidney function. Previously unknown morphometric signatures of podocyte depletion were identified in patients with ANCA-GN, which allowed patient classification and showed potential for risk stratification in combination with routine clinical tools. Together, our approach enables robust and scalable molecular morphometric analysis of human tissues, yielding deeper biological insights into the human kidney pathophysiology.

**Summary:** Deep learning enables robust and scalable molecular morphometric analysis of human tissues, yielding deeper biological insights into the human kidney pathophysiology.

## Introduction

The kidney continuously filters blood and maintains overall body homeostasis relying on a delicate balance between a complex vascular network and multiple specialized cell types^1^. Podocytes are kidney epithelial cells with limited capacity for regeneration that function as master regulators of glomerular health^2^. Experimental models show that severe podocyte loss leads to an irreversible process of progressive scarring rendering the affected glomeruli non-functional^3–5^. Furthermore, human podocyte loss has been identified in association with all major diseases contributing to chronic kidney disease^6–13^.

Anti-neutrophil cytoplasmic antibody associated glomerulonephritis (ANCA-GN) is primarily a systemic vasculitis with a strong immune-mediated epithelial reaction in the kidney, which leads to the formation of destructive glomerular lesions and a rapid loss of kidney functions^14^. While ANCA-GN has well-defined cellular changes^15^ that include podocyte injury^16^, podocyte loss is yet to be characterized in ANCA-GN patients. Using indirect immunofluorescence imaging, it is now possible to visualize different podocyte structures, facilitating the unambiguous identification of podocytes and thereby quantification of podocyte depletion^5,6^. However, reliable image segmentation for routine clinical analysis remains challenging, mostly due to time-constraints for detailed quantitative analysis with cellular resolution and lack of accuracy in available automated methods.

Time-constraints, precision and reproducibility are known hurdles in histopathology. For this reason, automation of classification and quantification processes has the potential to lessen the diagnostic burden and improve the quality of acquired data. Deep learning is increasingly gaining attention in multiple biomedical areas due to its potential clinical applications^17^, including natural language processing (i.e. analysis of electronic health records), and computer vision (i.e. histopathology and radiology). U-Net-based frameworks^18,19^ are particularly interesting for histopathology as they can be used for image segmentation and specific tasks such as image-to-image translation^20,21^. To date, multiple reports have shown the high-performance of deep learning networks for tissue-based classification of human disease^22,23^, but their role in detailed cellular morphometric profiling of clinical tissues remains unclear.

In this study, we present a deep learning-based workflow to perform cell-specific morphometric profiling of human kidney biopsies, including numbers, sizes, densities and distributions of podocytes within their respective glomerulus, which allowed a comprehensive characterization of endpoint variability within and between patients. We analyzed a total of 1095 glomeruli from 110 patients to profile 27,696 podocytes based on tissue expression of two complementary antigens in order to identify, segment and quantify podocyte depletion. A previously unrecognized morphometric signature of podocyte depletion was detected in patients with ANCA-GN, allowing patient classification with close to human accuracy and showing potential for risk stratification in combination with established clinical tools. Unexpectedly, we identified focal podocyte loss as a transitional state before the onset of overt lesion formation in patients with ANCA-GN, suggesting that podocyte injury may play a direct role in the pathophysiology of ANCA-GN. Together, these findings highlight the potential for deep learning-based architectures to enable robust and scalable molecular morphometric analysis of human tissues.

## Results

### Morphometric profiling of human samples using a dual segmentation U-Net

Human kidney biopsies from patients with available clinical data (i.e. age, sex, and eGFR), pathological endpoints (i.e. interstitial fibrosis) and integrative scores (i.e. ANCA-GN score) were immunolabeled using antibodies against podocyte-specific transcription factors, including nuclear expression of Dachshund Family Transcription Factor 1 (DACH1) and cytoplasmic expression of Wilms’ Tumor 1 (WT1), in order to unambiguously identify glomerular podocytes (**Fig. 1a**) and carefully profile a total of 27,696 podocytes. 1095 immunolabeled images were used for training, validation and testing during the development of the deep learning architectures (**Table S1**), including 722 images from 48 controls and 373 images from 62 patients with ANCA-GN. General patient demographics are outlined in **Fig. S1**.

**Figure 1.**
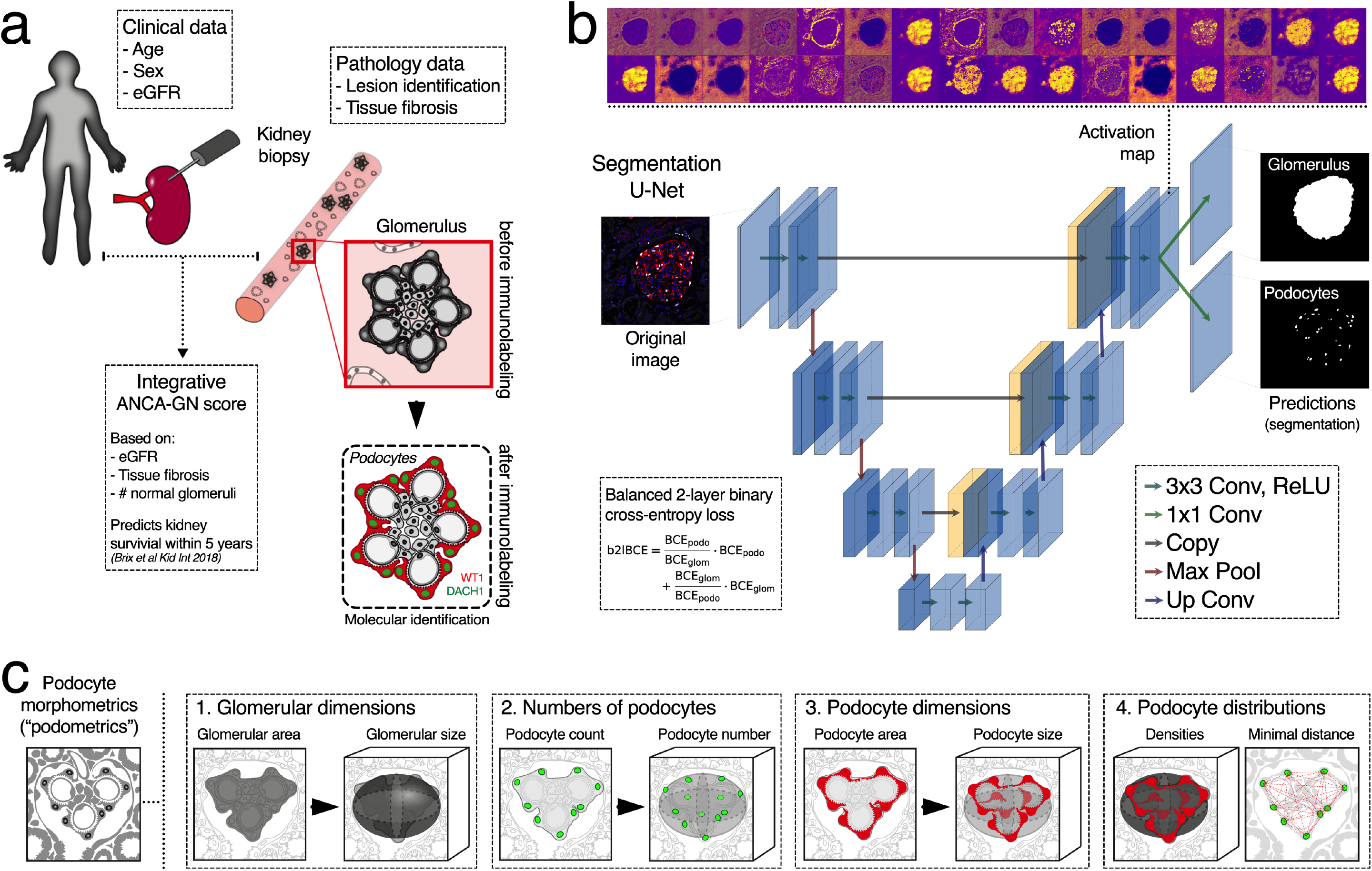
Segmentation U-Net for molecular morphometrics in kidney samples. (a) Biopsies from patients with immune-mediated kidney diseases, which are diagnosed, treated and monitored based on clinical, pathological, and integrative data, are used to perform molecular labeling of kidney podocytes, based on indirect immunofluorescence. (b) Glomerular area, and podocyte nuclei are virtually dissected from high-resolution confocal images with a segmentation U-Net for two simultaneous outputs that was trained using a balanced 2-layer binary cross-entropy loss. (c) 3D podocyte morphometrics (podometrics) were generated by model-based stereology, which extrapolates 3D from 2D data; in this case, glomerular and podocyte areas, and podocyte spatial location were used to estimate 3D glomerular dimensions, and numbers, sizes and distributions of podocytes. ANCA-GN: Anti-neutrophil cytoplasmic antibody associated glomerulonephritis; DACH1: Dachshund Family Transcription Factor 1; WT1: Wilms’ Tumor 1; BCE: binary cross-entropy; and Conv: convolution.

We developed a dual output segmentation U-Net that has an encoder/decoder structure with three convolutional layers, each containing between 32 (first/last layers) and 256 filters (bottom layer), which can simultaneously extract glomerular and podocyte nuclear areas from a composite fluorescent image (**Fig. 1b**). Segmented areas are integrated into model-based stereology formulas that estimate podocyte morphometrics (podometrics), including glomerular dimensions, numbers of podocytes, and podocyte dimensions and distributions (i.e. minimal distances between neighboring podocytes) within each glomerulus (**Fig. 1c**).

In order to secure optimal performance of the U-Net, hyperparameters were determined in a cross validation, where we confirmed that the number of training images was sufficient to achieve Dice scores over 0.90 (**Fig. S2a**). Network training was controlled via a balanced 2-layer binary cross-entropy loss that adaptively accounts for the performance of each segmentation task (**Fig. S2b-d**). Our dual output segmentation U-Net was compared to two single output U-Nets (for glomerular and podocyte nuclear areas separately), showing similar results (**Fig. S3a-b**). Furthermore, our dual output U-Net outperformed a customized ImageJ-based segmentation script at pixel and object levels, with a strong reduction on false positive rates (**Fig. S4a-b**).

### U-Net cycleGAN for annotation-free bias minimization

A lack of generalization is a well-known vulnerability of deep learning architectures^17^. To this end, we first compared podometrics obtained from the same patients that were systematically imaged in two different locations with different microscopes and by different operators with different levels of microscopy experience. Podocyte density was not affected by these different conditions (**Fig. S5a**), neither at a patient-level nor at a glomerular level (**Fig. S5b**), when the segmentation U-Net was trained jointly on these datasets. However, using the variance in DACH1 or WT1 expression per image (pixel level), significant differences were observed (**Fig. S5c**), suggesting that batch effects and image bias should be addressed in order to increase the reproducibility and scalability of the method.

Multiple operators and microscopes lead to differences in image quality, differing from the reference dataset (**Fig. 2a**). One solution is to continuously re-train the segmentation U-Net (**Fig. 2b**), which progressively leads to a more robust network, but requires manual annotations. An alternative approach can be found in the use of deep learning-based annotation-free bias minimization (**Fig. 2c**). Thus, we implemented a U-Net cycleGAN (cycle-consistent generative adversarial network with a U-Net like generator) to transform images obtained under different conditions (i.e. microscope and operator) into images resembling the reference dataset used for training the segmentation network (**Fig. 2d**). Representative images show the resulting segmentation optimization (**Fig. 2e**) and improvements in Dice scores at both pixel and object levels (**Fig. 2f**). Training curves of the U-Net cycleGAN, as well as ROC and precision-recall curves for the different combinations of data and segmentation networks, are provided in **Fig. S6a-b**. While these results provide evidence that unannotated datasets can be efficiently segmented using a network trained on the reference dataset when they are bias transferred using the U-Net cycleGAN (e.g. improvement of the mean podocyte pixel-based Dice score from 0.65 to 0.81, see center and right panel of **Fig. S6c**), we obtained slightly better segmentation results using a segmentation U-Net trained on all manually segmented data, including the two control and the ANCA-GN datasets (mean podocyte pixel-based Dice score: 0.84, left panel of **Fig. S6c**). All further results are therefore based on the U-Net trained jointly on the two control and the ANCA-GN datasets.

**Figure 2.**
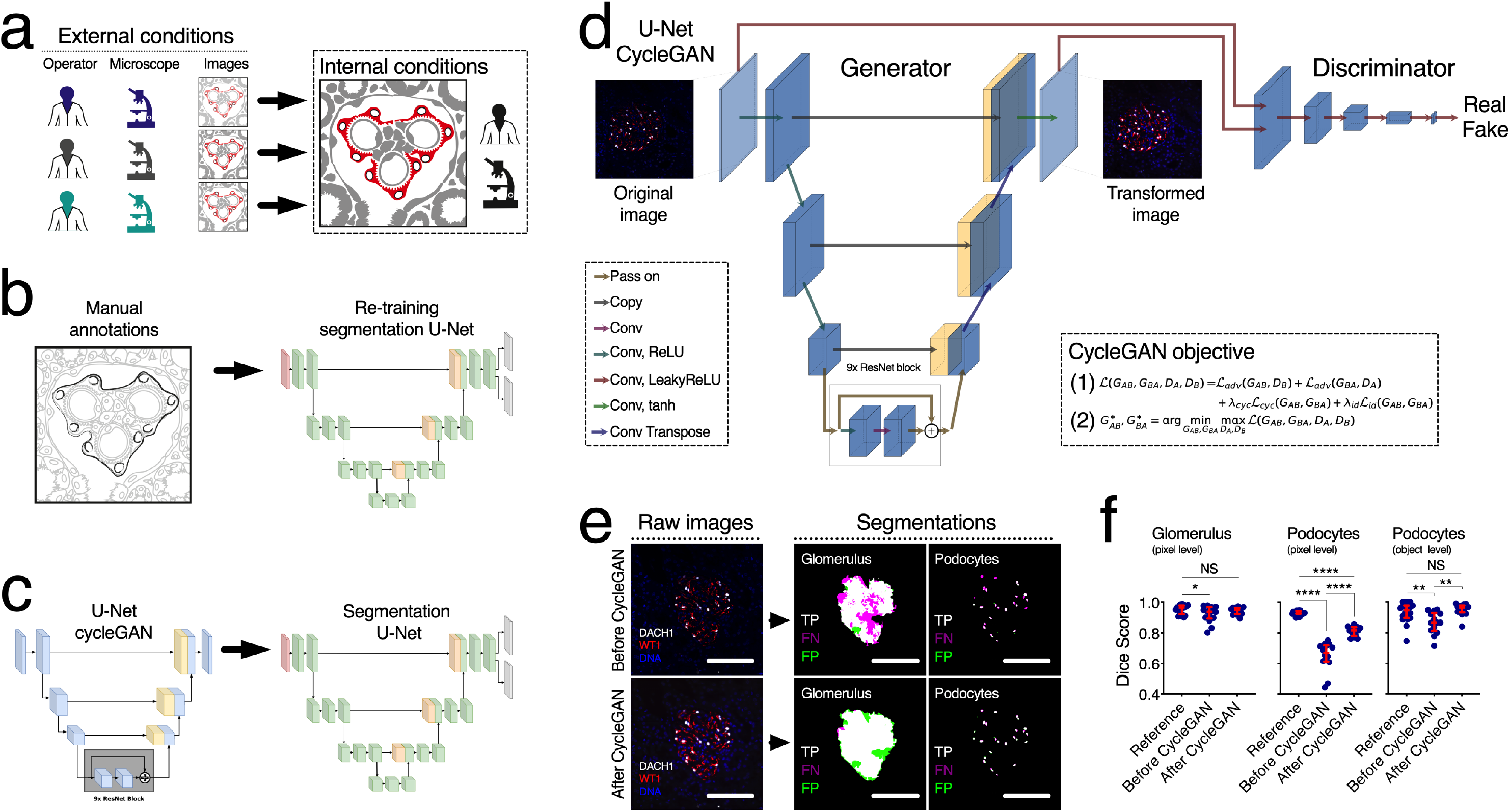
U-Net cycleGAN for bias minimization between different dataset domains. (a) Scalable frameworks require adaptability to external conditions. While indirect immunofluorescence protocols can be standardised, operator training, microscopy set-up and eventually image quality are hard to control, especially if segmentation tasks have been defined based on tightly controlled internal conditions. (b) We propose using a U-Net cycleGAN (without annotations) in order to transform images before applying the segmentation U-Net. (c) Alternatively, manual annotations can be performed in order to re-train the segmentation U-Net before it is applied to a new dataset. (d) While the generator in the U-Net cycleGAN transforms images from one dataset domain to the other, the discriminator tries to distinguish between ‘real’ and ‘fake’ images. This adversarial game is reflected in the cycleGAN objective, which is made up of the adversarial loss L_adv_, the cycle-consistency loss L_cyc_ and an identity loss L_id_. (e) Representative images showing segmentation agreement with ground truth and reductions in false negatives. (f) Dice score at both pixel and object-level significantly improved after cycleGAN for podocytes. In dot plots, every blue dot represents one image and red error bars represent medians and interquartile ranges. TP: true positives, FP: false positives, FN: false negatives. ****P<0.0001, **P<0.01, *P<0.05, and NS: not statistically significant. Scale bars 150μm.

### Molecular podometrics reveal podocyte loss in ANCA-GN

Representative images show high accuracy and precision of the U-Net for podocyte segmentation in samples from both controls and ANCA-GN patients (**Fig. 3a**), which was illustrated by receiver operating characteristic (ROC) and precision-recall curves (**Fig. 3b**). Strong agreement between ground truth and U-Net outputs was determined by pixel- and object-based Dice scores (mean podocyte Dice scores for controls 0.86 and 0.95 respectively) (**Fig. 3c**). While image segmentation in ANCA-GN patients was comparable to controls, detection levels were not identical (mean podocyte Dice scores for ANCA-GN patients 0.87 and 0.91 respectively). For this reason, we compared segmented areas obtained as ground truth and from the U-Net, which showed identical differences between controls and ANCA-GN patients (**Fig. S7a-b**), which supports biological differences rather than technical artefacts. Furthermore, we also determined direct correlations between ground truth and U-Net segmentation outputs from both controls and ANCA-GN patients (**Fig. S7c**).

**Figure 3.**
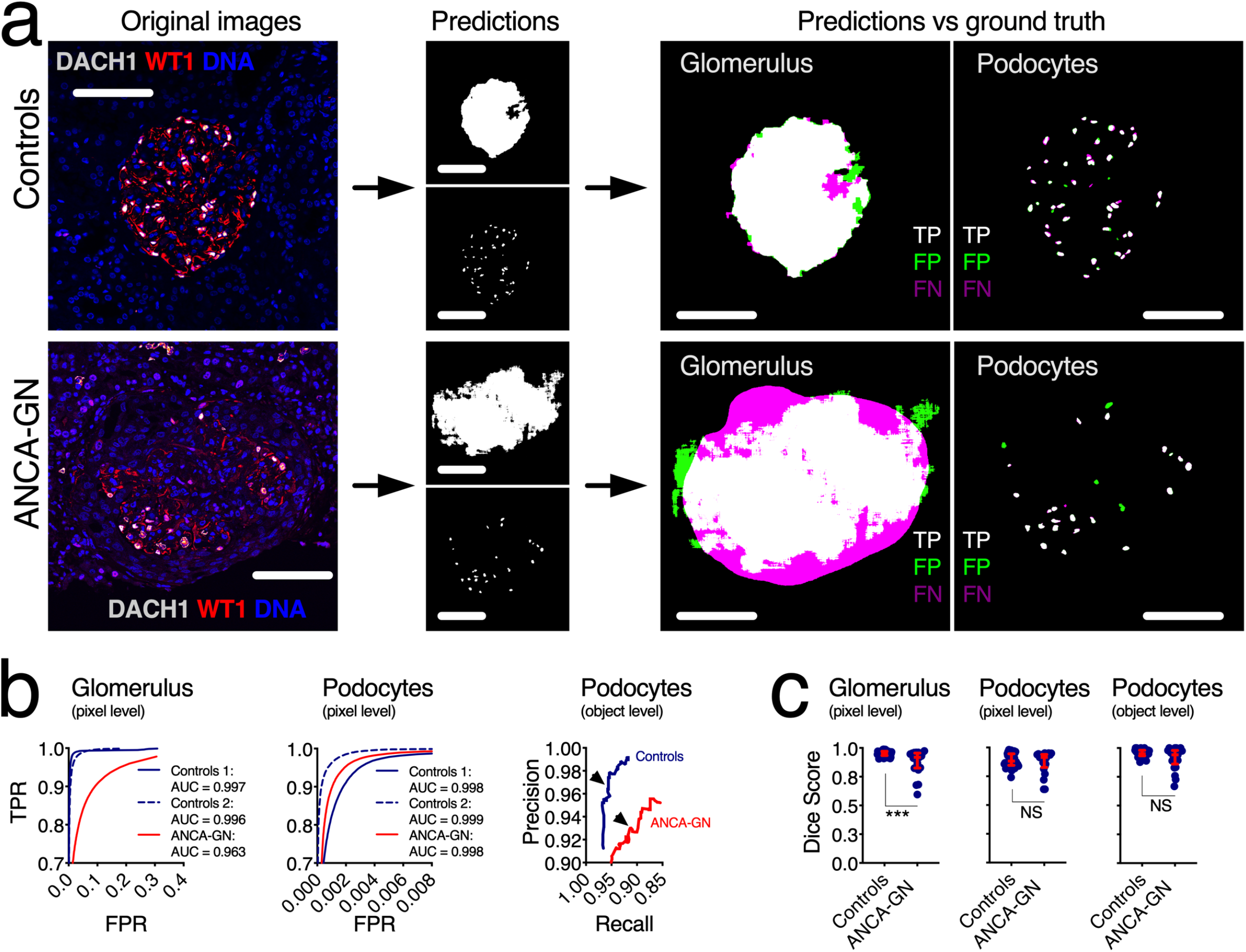
Application of segmentation U-Net to human kidney biopsies. (a) Visual representation of the segmentation process, from original images, to segmentation outputs for glomeruli and podocytes, and their respective correlation with manually-segmented ground truths, highlighting true positives, false positives and false negatives. (b) Receiver operating characteristic (ROC) and precision-recall curves in samples from controls and ANCA-GN patients – arrowheads show selected thresholds for both conditions. (c) Dice scores at pixel and object levels for glomeruli and podocytes, showing comparable segmentation performance in health and disease. In dot plots, each blue dot represents one image, red error bars represent medians and interquartile ranges. TPR: true positive rate, FPR: false positive rate, AUC: area under the curve, TP: true positives, FP: false positives, FN: false negatives. ***P<0.001 and NS: not statistically significant. Scale bars in all panels represent 100μm.

Reductions in median podocyte numbers and densities with consequent increases in median podocyte sizes and distances between closest neighbors were found in patients with ANCA-GN compared to controls (**Fig. 4a**). Median glomerular size was directly associated with median podocyte number (R=0.48, P<0.0001 in controls and R=0.57, P<0.0001 in ANCA-GN) with significant differences in the intercept (P<0.0001), which suggests podocyte loss across the entire spectrum of glomerular size (**Fig. 4b**). Similarly, median podocyte density was inversely associated with median minimal distances between neighboring podocytes (R= −0.88, P<0.0001 in controls and R=0.68, P<0.0001 in ANCA-GN) with statistical differences in the slope (P<0.01), suggesting that compensatory podocyte hypertrophy is exacerbated in ANCA-GN patients (**Fig. 4c**).

**Figure 4.**
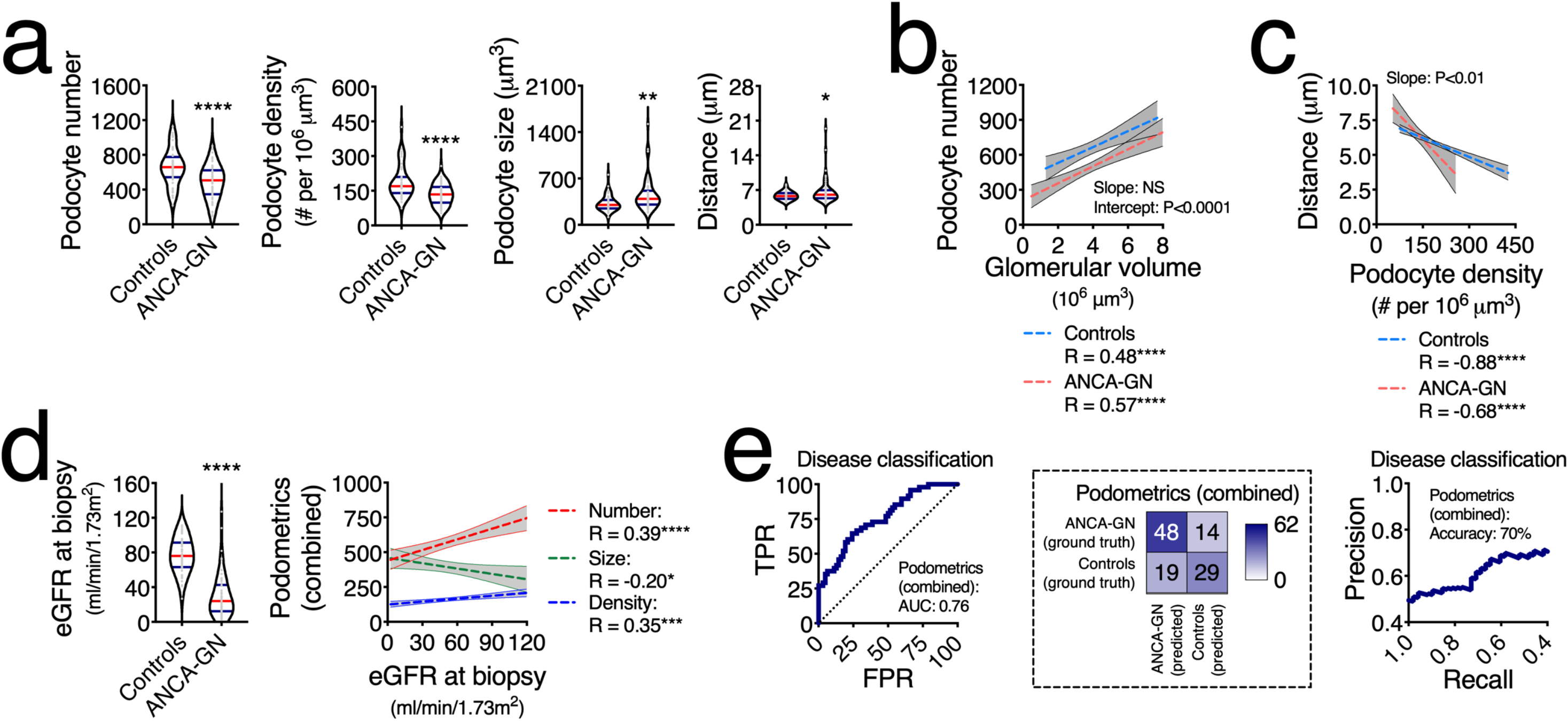
Molecular podometrics reveal podocyte loss in patients with ANCA-GN. (a) Podocyte morphometric analysis (podometrics; median per patient) showing reductions in podocyte numbers and densities, and increases in podocyte sizes and closest neighbour distances in ANCA-GN patients compared to controls. (b) Correlation analyses confirm a pattern of podocyte loss across the entire range of glomerular volume. (c) Increases in podocyte closest neighbour distances are associated with reductions in podocyte density. (d) ANCA-GN patients have lower estimated glomerular filtration rate (eGFR) at the time of biopsy compared to controls – features of podocyte depletion are associated with eGFR at biopsy. (e) Receiver operating characteristic (ROC) and precision-recall curves showing discrimination power of combined podometrics (podocyte number, density and size), including confusion matrix. In violin plots, each grey dot represents the median value per subject, red lines represent medians and blue lines interquartile ranges. Regression lines represent lines of best fit and 95% confidence intervals. TPR: true positive rate, FPR: false positive rate, AUC: area under the curve. ****P<0.0001, **P<0.01, and *P<0.05.

In our cohort, the main clinical discriminator between controls and ANCA-GN patients was kidney function, assessed by estimated glomerular filtration rate (eGFR) at the time of biopsy. In particular, eGFR was associated with podocyte number (R=0.39, P<0.0001), density (R=0.35, P<0.001) and size (R= −0.20, P<0.05) (**Fig. 4d**). Using a leave-one-out cross-validation approach, we generated a combined podometric score, including podocyte number, density and size, which also partially discriminated between controls and ANCA-GN patients with an AUC of 0.76 (**Fig. 4e**). Together, these findings suggest a potential overlap in the levels of podocyte depletion between controls and ANCA-GN patients.

### Morphometric signature of podocyte depletion identifies patients with ANCA-GN

Lesion development in ANCA-GN is focal, meaning that within the same patient some glomeruli are affected, and others are not (**Fig. S8a**). This is directly reflected in the changes in the variances per subject of all podometric parameters (**Fig. S8b**), which decreased in podocyte numbers and densities, but increased in sizes and distances between closest neighbors.

Analyses of single glomeruli showed that podocyte depletion was present in ANCA-GN patients, even in glomeruli that were not defined as glomerular lesions and was associated with compensatory podocyte hypertrophy (**Fig. 5a**). Principal component analysis also revealed that normal glomeruli in ANCA-GN patients represent a transitional state from normal glomeruli in controls to overt glomerular lesions in ANCA-GN (**Fig. 5b**), suggesting that analyses of individual glomeruli within one patient may provide additional clues that may be applied to differentiate controls and ANCA-GN patients. Using a leave-one-out cross-validation approach, we generated a morphometric signature of podocyte depletion, which is generated per subject based on all available morphometric data, including both central tendencies and measures of variability. Importantly, this integrative parameter discriminated between controls and ANCA-GN patients as shown in both ROC (**Fig. 5c**) and precision-recall curves (AUC: 0.88) with an accuracy of 82% (**Fig. 5d**), which was almost identical to the discrimination power of eGFR (AUC: 0.92 and accuracy: 86%).

**Figure 5.**
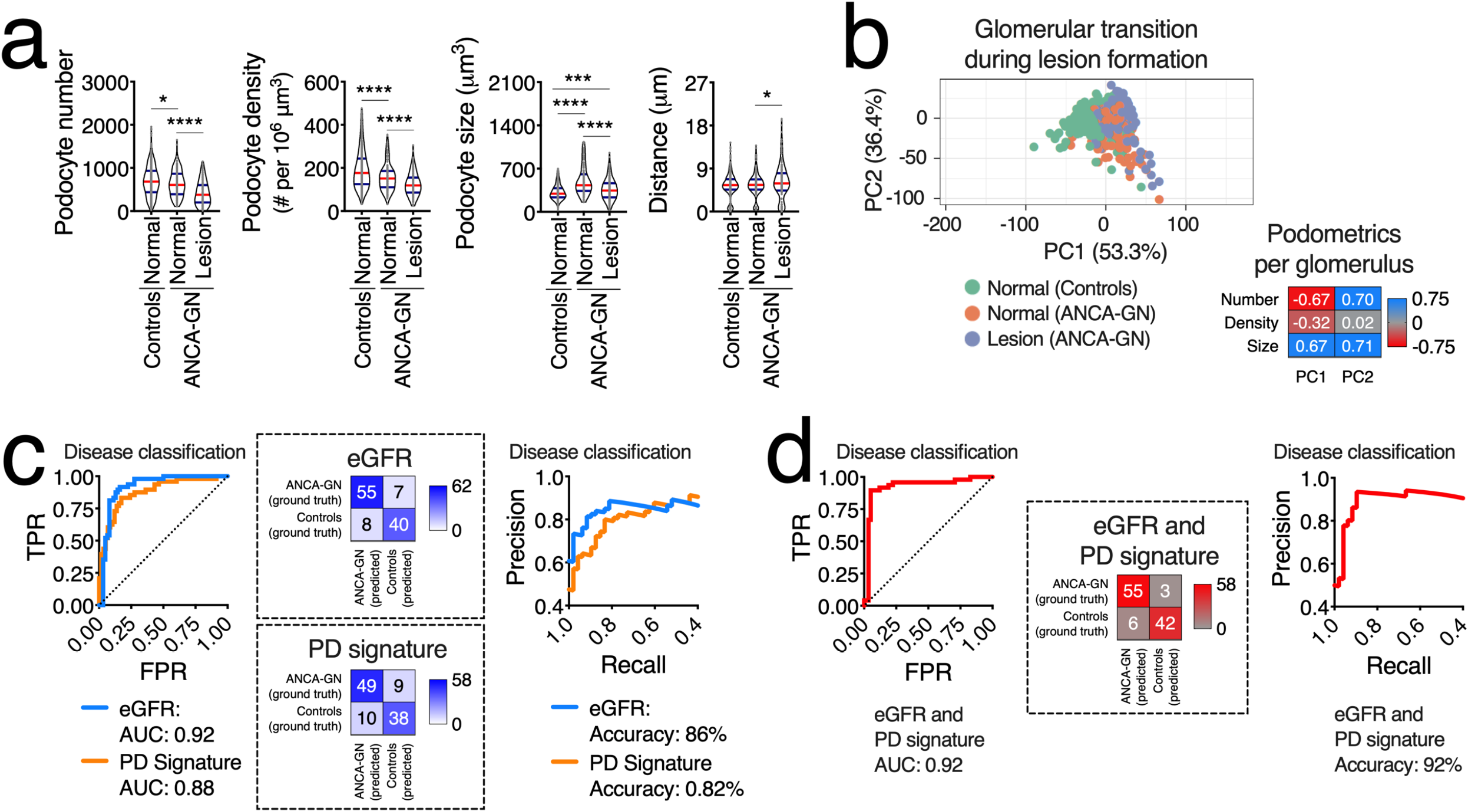
Podocyte morphometric signature identifies ANCA-GN patients. (a) Podocyte morphometric analysis (podometrics; per glomerulus) showing a pattern of podocyte loss and hypertrophy in glomeruli classified as “normal” in ANCA-GN patients. (b) Principal component (PC) analysis using Pareto scaling to rows. Probabilistic PCA was used to calculate principal components, confirming that normal glomeruli in ANCA-GN patients represent a transitional state between normal glomeruli in controls and lesions in ANCA-GN patients. (c) Receiver operating characteristic (ROC), precision-recall curves and confusion matrices of patient classification based on eGFR and on a morphometric signature of podocyte depletion (PD), which combines morphometric data from every available glomerulus per biopsy per patient. (d) ROC, precision-recall curves and confusion matrices for eGFR and PD as classifiers. In violin plots, each grey dot represents one glomerulus, red lines represent medians and blue lines interquartile ranges.. TPR: true positive rate, FPR: false positive rate, AUC: area under the curve. ****P<0.0001, ***P<0.001, and *P<0.05.

In this cohort, 3 ANCA-GN patients were classified as controls and 6 controls were classified as ANCA-GN. First, we hypothesized that this could be due to segmentation artefacts as DACH1 expression is upregulated in other cell types (i.e. erythrocytes and proximal tubular cells). We carefully screened all images from these 9 misclassified subjects and confirmed appropriate segmentation – representative images are shown in **Fig. S9a**. In patients with ANCA-GN, misclassified subjects were younger and had higher eGFR than median values for controls. In controls, misclassified cases were older and had lower eGFR than median values for ANCA-GN patients (**Fig. S9b-c**). Together, these findings suggest that misclassifications may be associated with early stages of disease in ANCA-GN and age-related podocyte loss in controls. Furthermore, this morphometric signature of podocyte depletion marks the degree of disease progression in close relation to physiological readouts.

### Potential of podometrics for risk stratification in patients with ANCA-GN

A recent study proposed an integrative predictive score of 5-year kidney survival in ANCA-GN^24^, based on eGFR, percentage of interstitial fibrosis and number of “normal” glomeruli. We adapted this ANCA score to include a baseline comparison to control patients and model associations to podometrics, showing that median podocyte number, density and size are significantly correlated with the modified ANCA-GN score (**Fig. 6a**). From a total of 62 patients with ANCA-GN, 58 had at least 3 identified glomeruli in the diagnostic biopsy, which allowed us to perform analysis of intra-subject variability. From these subjects, 8 showed a poor outcome, defined as mortality, relapse or loss of at least 10% of eGFR within their respective follow-up period (**Fig. 6b**). For a balanced comparison, we carefully age and sex matched these 8 subjects within remaining available patients from our cohort (n=8 matched ANCA-GN patients). While our matching strategy was successful for age and sex, we were not able to obtain matches by eGFR (**Fig. 6c**). Variances (**Fig. 6d**) and ranges (**Fig. 6e**) in podocyte size were significantly increased in ANCA-GN patients with poor outcomes. Neither the conventional ANCA-GN score nor our adapted version were different between the outcome groups (**Fig. 6f**), but a ratio between the adapted ANCA-GN score and range in podocyte size showed significant differences by outcome group (**Fig. 6g**). In summary, these findings highlight a potential for risk stratification among ANCA-GN patients using a combination of podometrics and clinical tools.

**Figure 6.**
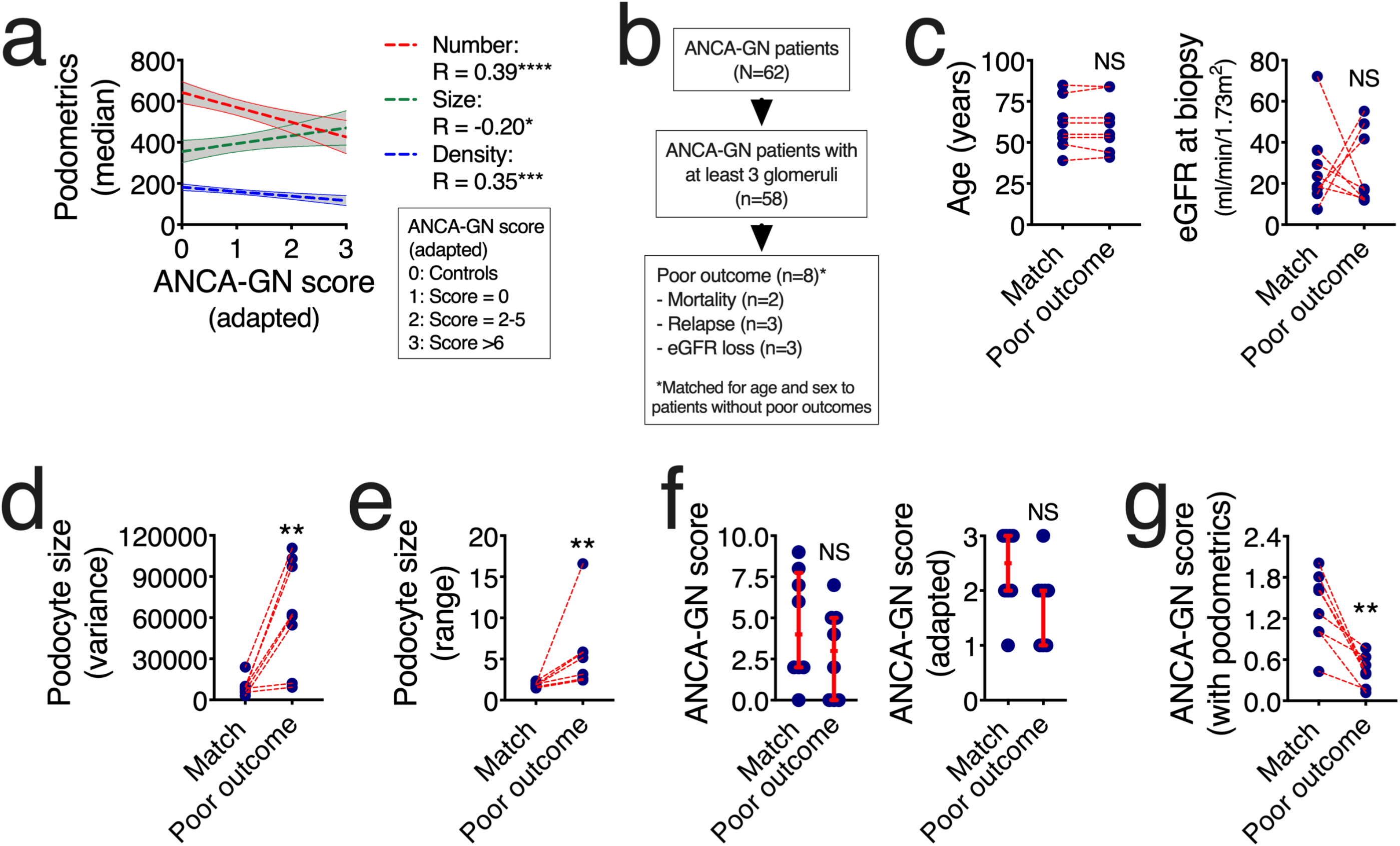
Potential role of podometrics for ANCA-GN risk evaluation. (a) Features of podocyte depletion correlate with an adapted ANCA-GN score that predicts poor clinical outcomes within 5 years. (b) Among all 62 ANCA-GN patients, clinical follow-up data identified a total of 8 patients with poor clinical outcomes, including mortality, relapse and loss of estimated glomerular filtration rate (eGFR) of at least 15% from baseline, which were carefully age and sex-matched to patients without negative outcomes. (c) Successful age-match with random selection of variable eGFR. (d) Variance in podocyte size per biopsy was significantly elevated in patients with poor outcomes. (e) The ratio between maximal and minimal podocytes sizes (range) per biopsy was also increased in patients with poor outcome. (f) Neither the classical ANCA-GN score nor an adapted ANCA-GN score are different between patients with poor outcome and matched controls. (g) A modified ANCA-GN score based on a ratio between the adapted ANCA-GN score and the range of podocyte size per biopsy was significantly reduced in patients with poor outcome. Regression lines represent lines of best fit and 95% confidence intervals. Each blue dot represents one subject. In (f) red lines represent medians and interquartile ranges. ****P<0.0001, ***P<0.001, **P<0.01, *P<0.05 and NS: not statistically significant.

## Discussion

In this study, we present a deep learning-based approach that automatically identifies morphometric signatures of podocyte depletion in human kidney biopsies, achieving human-level accuracy while saving time and resources. Our method provides robust and scalable molecular morphometric endpoints for patients with ANCA-GN, revealing novel pathophysiological insights of kidney epithelial biology and serving as an example for successful integration of deep-learning-based technologies into clinical settings.

Previous deep learning studies focused on the end-to-end evaluation of biopsies through classification into several categories based on classical histology^22,23,25–27^. To our knowledge, this is the first report to integrate deep learning for object segmentation in clinical samples that can be used for cell-specific morphometrics, which not only allows disease classification and risk stratification, but also provides objective endpoints for the analysis of kidney biopsies. Furthermore, antigen-based cellular identification reduces subjectivity in annotation strategies, as extensive specialized training is not needed in order to identify protein expression with fluorescence microscopy, accelerating annotations and homogenizing ground truth definition, all of which are well-defined obstacles for clinical translation of deep learning-based methodologies^17^. However, reproducibility remains a valid drawback for new clinical tools, especially those dependent on microscopy.

Bias minimization through a U-Net cycleGAN allows a wider use of the pipeline given that data obtained by various users and on different microscopes can be adapted in order to efficiently homogenize image quality. Generative networks have been used in the past for histopathological analysis, but mostly have been limited to classical histological stainings^28,29^. While this strategy is certainly effective and is comparable to multi-dataset training, manual annotations and re-training of the segmentation architecture is the safest approach to maximize accuracy. In this manuscript, we provide both options, allowing users to decide based on their experimental and clinical needs.

The limited sample size for training, optimization, validation and testing of multiple deep learning architectures may be perceived as a shortcoming of the present study. However, this is a very common problem in biomedical sciences. The number of patients with follow-up data and negative outcomes prevented us to provide predictive analyses at this stage. For this reason, our observations should be carefully validated in larger cohorts with longer follow-up periods. Furthermore, the successful integration of artificial intelligence-based morphometrics into clinical practice will not only depend on larger datasets but also on standardization and automation of tissue processing and imaging. While our efforts for batch effect minimization are promising, we only tested variations in image quality based on two parameters: microscopy operators and confocal systems. The compatibility of our approach with other high-throughput imaging methodologies, such as spinning disk and epi-fluorescence-based systems, still needs to be validated.

The devastating nature of ANCA-GN requires continuous efforts to identify diagnostic and prognostic tools that may guide clinical management^15^. In the pathophysiology of lesion formation during ANCA-GN development and progression, it is known that immune cells and parietal epithelial cells play key roles^1,14,16^. Importantly, our data highlight a previously unrecognized role of podocyte loss in ANCA-GN that could only be revealed by analyzing single glomeruli and their variability within and between subjects. The unexpected value of podometric endpoints in diagnostic ANCA-GN biopsies can only strengthen the position of podocyte depletion as a hallmark of glomerular disease^30,31^. Future studies will assess whether podocyte depletion signatures may serve as objective endpoints for the management of glomerular diseases, and their potential applicability to patient diagnosis, management and prognosis. It is our hope that this study may pave the way into the development and implementation of advanced tissue morphometrics in routine clinical pathology.

## Author contributions

Study initiation, TBH, SB and VGP; Conceptualization, MK, MZ, TBH, SB and VGP; Methodology and analysis, MZ, MK, MNW, AKT, LG, CK, MH, JK, NW, FB, SW, TW, UP, CFK, EH, RK, SB and VGP; Writing, MZ, MK, AKT, TBH, SB, and VGP; Supervision, TBH, SB and VGP.

## Data and materials availability

The datasets generated and analyzed during the current study are available from the corresponding authors on reasonable request. Codes will be made available via GitHub upon publication of the manuscript.

## Materials and Methods

### Human tissue

Tissue collection from nephrectomy samples due to renal cell carcinoma was performed at Eschweiler Medical Center (Germany). Ethics approval was obtained from the Institutional Review Board of the RWTH Aachen University Medical Center, Germany (EK-016/17). After fixation with 4% paraformaldehyde (PFA), representative kidney blocks from the pole opposite to the tumor were extracted – a strategy that aimed to collect non-pathological tissue. Kidney biopsies from patients with ANCA-associated glomerulonephritis were obtained from the Hamburger Glomerulonephritis Registry (ethics approval: PV4806).

### Immunofluorescence and confocal microscopy

Previously reported protocols were applied^5,6^. To identify podocytes, we used a combination of WT1 (Agilent Technologies; IS05530-2) and DACH1 (Sigma-Aldrich; HPA012672)^32^ as primary antibodies, Alexa-Fluor 488, 555 and/or 647 as secondary antibodies (Invitrogen; A21202, A31572, A31571, and A31573) depending on the experiment, and a DNA marker to identify single nuclei, either DAPI (Sigma-Aldrich; D9542) or DRAQ5 (Abcam; ab108410). Optical images were obtained using inverted laser confocal microscopes (Nikon and LSM800, Zeiss), stored in 1024×1024-pixel frames. Each image contained one glomerulus.

### Manual image annotation for ground truth generation

Ground truth datasets were generated based on podocyte nuclei and glomerular areas in manual segmentation performed by three expert scientists trained under equal conditions within our team, blinded from the patient data. Quality control was performed by a senior scientist within our team. During training, the segmentation U-Net then learned from the annotated images (training and validation sets) to segment the structures of interest and the final results were validated on another set of annotated images (test set).

Glomeruli were classified as normal or lesion based on anatomical criteria. Normal glomeruli had a monolayer of parietal epithelial cells and glomerular tufts with homogenous and robust podocyte labels, namely cytoplasmic WT1 and nuclear DACH1. Glomerular lesions showed at least a double layer of parietal epithelial cells and/or capillary collapse and/or segmental or global absence of podocyte labeling.

### ImageJ baseline script for glomerulus and podocyte nuclei segmentation

In order to segment the glomerulus using ImageJ, the following sequence was used: (1) channel splitting, (2) thresholding and then dilation applied to each channel separately, (3) channel merging, (4) filling holes, eroding, and particle analysis, and (5) selection of the biggest region of interest. In order to segment podocyte nuclei using ImageJ, the following sequence was used: (1) channel splitting and thresholding, (2) dilation of WT1 channel followed by combination of all channels using the logical operator “AND”, (3) thresholding of DNA label, (4) dilation, filling holes and eroding, and (5) distance transformation using watershed (MorphoLibJ plugin).

### Dual output segmentation U-Net

Inspired by Ronneberger et al.^1^ a U-Net architecture was implemented in Python 3 using Tensorflow 1.13. The segmentation U-Net consists of an encoder with three layers, where the convolutions in the first layer have 32 filters. The number of filters is doubled in the following layers. After the bottom layer with 256 filters, in each of the three layers of the decoder the number of filters is halved again. We pad the images in order to receive segmentations of the same size as the input images.

The U-Net was modified to simultaneously return a dual segmentation output: glomerular areas and podocyte nuclear areas. An annotated subset of images (n=317) was split into training (192 images), validation (60 images) and test (65 images) subsets with the relation of approximately 60/20/20, maintaining that all images from one subject should belong to one subset. For training, the use of extensive on-the-fly data augmentation (horizontal and vertical flips, horizontal and vertical shifts, rotations up to 45°) is important for the generalizability of the network. The network was trained for 2000 epochs with a batch size of 2 images on an Nvidia Tesla V100 graphics card. We used RMSprop as an optimizer and introduced a custom balanced two-layer binary cross-entropy loss that adaptively takes into account the current performance of each segmentation task. The binary cross-entropy for each task:

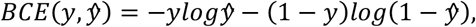

for mask *y* and prediction 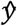 (the prediction 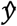 is clipped to lie between *ε* and 1 – *ε*, with *ε* = 1e-07, in order to avoid logarithms of 1 and thus later divisions by 0) is adapted to consider both segmentation tasks simultaneously and to weight each term so that the currently poorer performing task receives more importance:

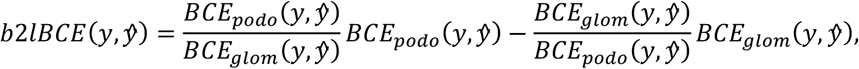

where *BCE_podo_* and *BCE_glom_* are the binary cross-entropies for the podocyte and glomerulus segmentation tasks respectively. Additionally, we weighted foreground objects and the background in the training loss in order to enforce a better segmentation of narrowly spaced podocytes. This was done using weight maps for each image similar to those proposed by Falk et al.^19^:

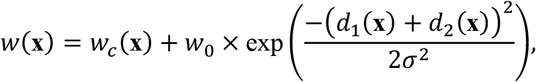

where *w_c_* is the class probability map for the mask, *d*_1_ is the distance to the border of the nearest cell and *d*_2_ is the distance to the border of the second nearest cell.

Our evaluation metric is the commonly used Dice score, evaluated for each task separately at pixel level and additionally at object level for the podocytes. Incomplete nuclear parts or glomeruli were filtered out using post-processing, removing all objects smaller than 800μm for glomeruli and 3μm for podocyte nuclei. Given that this architecture provided excellent results, and some tests with a more complex architecture (i.e. Mask R-CNN) yielded similar results, we decided to work with our more compact and faster dual U-Net.

### Hyperparameter optimization

In order to find the optimal architecture and hyperparameters, an extensive grid search across various options was performed using Ray Tune (https://ray.readthedocs.io/en/ray-0.3.1/tune) and Sacred (https://sacred.readthedocs.io/en/stable/) – chosen values in brackets: single vs. dual segmentation (dual), number of layers (3), number of filters in first layer (32), dropout in encoder and decoder (no), dropout in the bottom layer (yes), skip connections between encoder and decoder (yes), dropout in the skip connections (no), batch normalization (yes), optimizer (RMSprop), learning rate (1e-05), learning rate decay (no), loss (balanced two-layer binary cross-entropy), weighting (yes), histogram equalization (no), contrast stretching (no), data augmentation (yes), oversampling of crescents (no). To evaluate these, iteratively, a few (related) hyperparameters were varied. Then, using 4-fold cross validation on the combined training and validation subsets of the Controls 1 dataset, networks were trained and the optimal configuration was chosen based on the average Dice scores as well as their standard deviation between the different folds of the cross validation (or for similar performance, the least data/computationally intensive). This process was repeated with the next set of hyperparameters. In a similar fashion, using a 10-fold cross-validation, we evaluated the number of images used for training to ensure that approximately 60-70 images per dataset yielded satisfactory results with Dice scores above 0.90.

### U-Net cycleGAN configuration

The U-Net cycleGAN was implemented in Python 3 with Tensorflow 2.0. The generator is made up of an encoder, a transformer, and a decoder. Based on Zhu et al^20^ the encoder consists of three convolutional layers with 64, 128 and 256 filters, kernel sizes 7, 3 and 3, and strides 1, 2, and 2. All layers use instance normalization as well as ReLU activation. The transformer consists of 9 ResNet blocks^33^, which are made up of two convolutional layers with instance normalization and ReLU activation for the first layer. The decoder is made up of three transpose convolution layers. Before each, the input is concatenated with the output of the corresponding layer in the encoder. The transpose convolution layers have 128, 64 and 3 filters, kernel sizes 3, 3 and 7, and strides 2, 2, and 1. All use instance normalization, but the first two are ReLU activated unlike the last one, which has a Tanh activation since the output of the last layer is the generated image which should have pixel values between −1 and 1. The discriminator consists of 6 convolutional layers, with 64, 128, 256, 512, 512 and 1 filters, kernel size 4, and strides 2, 2, 2, 2, 1 and 1. Except for the first and last layer all are instance normalized. And, except for the last layer, all use a leaky ReLU activation with an alpha slope of 0.2.

### CycleGAN training

The model has been trained on 285 images from Controls 2 and 180 images from Controls 1. For the validation, 46 images from Controls 2 and 44 from Controls 1 have been used. The images are resized to 256×256 pixels with three channels (RGB) using Gaussian pyramids. After transformation, the images are upsampled to the original size of 1024×1024 pixels using Laplacian pyramids, as has been done in Engin et al^34^, consisting of layers calculated based on the original input. The network was trained for up to 200 epochs with a steady learning rate of 2e-04 for the first 100 epochs and a linearly decaying learning rate that ends at 0 after 200 epochs. The losses have been weighted with *λ_cyc_* = 10, and *λ_id_* = 5. The batch size was 1. The epoch with the lowest validation loss has been selected for translating the images (epoch 83). The network was trained on an NVIDIA Quadro RTX 8000 48GB with TeslaLink.

Since bias between different datasets is not a new problem, a comparison between generative models and traditional approaches was necessary. Since the (initial) effort for generative models is higher, it should be proven that they lead to better results. As baseline methods, histogram equalization, color transfer and an adaptation of the mean colors to the reference have been tested. However, none of these methods showed a substantial improvement.

### Molecular podometrics

Model-based stereology was applied to confocal images^35^ and allowed the estimation of podocyte number and podocyte density per glomerulus. Fiji imaging software (Max Planck Institute of Molecular Cell Biology and Genetics, Dresden, Germany) was used to navigate the raw files. Podocytes were defined as DAPI+WT-1+DACH1+ cells. Glomerular cross-sectional area was measured in order to estimate glomerular volume, and thereby define podocyte density.

The morphometric signature combines the podometrics per glomerulus within each patient by calculating the minimum, maximum, mean, median and variance of podocyte number, podocyte density, podocyte distance, podocyte nuclear area and glomerular area across all glomeruli per subject.

### Statistics

All statistical analyses were performed using GraphPad Prism (v8.0.2) and Stata 13.1. Results are reported as median and IQR. Significance was evaluated using the unpaired Mann Whitney’s test when comparing two continuous variables. For comparison of 3 groups, Kruskal-Wallis test with Dunn’s multiple comparisons test was used. Correlation analyses were performed using spearman rank coefficients. A p-value below 0.05 was considered to be statistically significant.

Classification of subjects into controls and ANCA-GN patients was performed in scikit-learn^36^ using a logistic regression and leave-one-out cross validation, where iteratively one subject was excluded from the training of the model and then used as test set. The final results are a combination of all subjects’ results, each thus tested on a different model. Due to the nature of leave-one-out cross validation without a completely unseen test set, no further optimization of parameters was possible. Each of the features in the morphometric signature was normalized by removing its mean and dividing by its standard deviation before using it to train the logistic regression (excluding the test subject). To evaluate podometrics at the level of single glomeruli, individual glomeruli were clustered based on podometrics via Principal Component Analysis (PCA) using Pareto scaling to rows. Probabilistic PCA was used to calculate principal components.

**Table S1.** Overview of all image datasets. This table summarises all used subjects and images, divided by training, validation, test and unannotated sets in both controls and ANCA-GN patients. ^$^denotes controls 1 and 2 shared 16 subjects for a total of 48 analysed controls.

| Dataset | Subset | # subjects | # images / glomeruli |
| --- | --- | --- | --- |
| Controls 1 | Training | 6 | 68 |
| Controls 1 | Validation | 2 | 21 |
| Controls 1 | Test | 2 | 20 |
| Controls 1 | Unannotated | 16 | 168 |
| Controls 1 | All | 26<sup>\$</sup> | 277 |
| Controls 2 | Training | 5 | 64 |
| Controls 2 | Validation | 2 | 20 |
| Controls 2 | Test | 2 | 24 |
| Controls 2 | Unannotated | 29 | 337 |
| Controls 2 | All | 38<sup>\$</sup> | 445 |
| ANCA-GN | Training | 10 | 60 |
| ANCA-GN | Validation | 4 | 19 |
| ANCA-GN | Test | 3 | 21 |
| ANCA-GN | Unannotated | 45 | 273 |
| ANCA-GN | All | 62 | 373 |
|  |  | # subjects | # images / glomeruli |
| **TOTAL** |  | 110 | 1095 |

**Figure S1.**
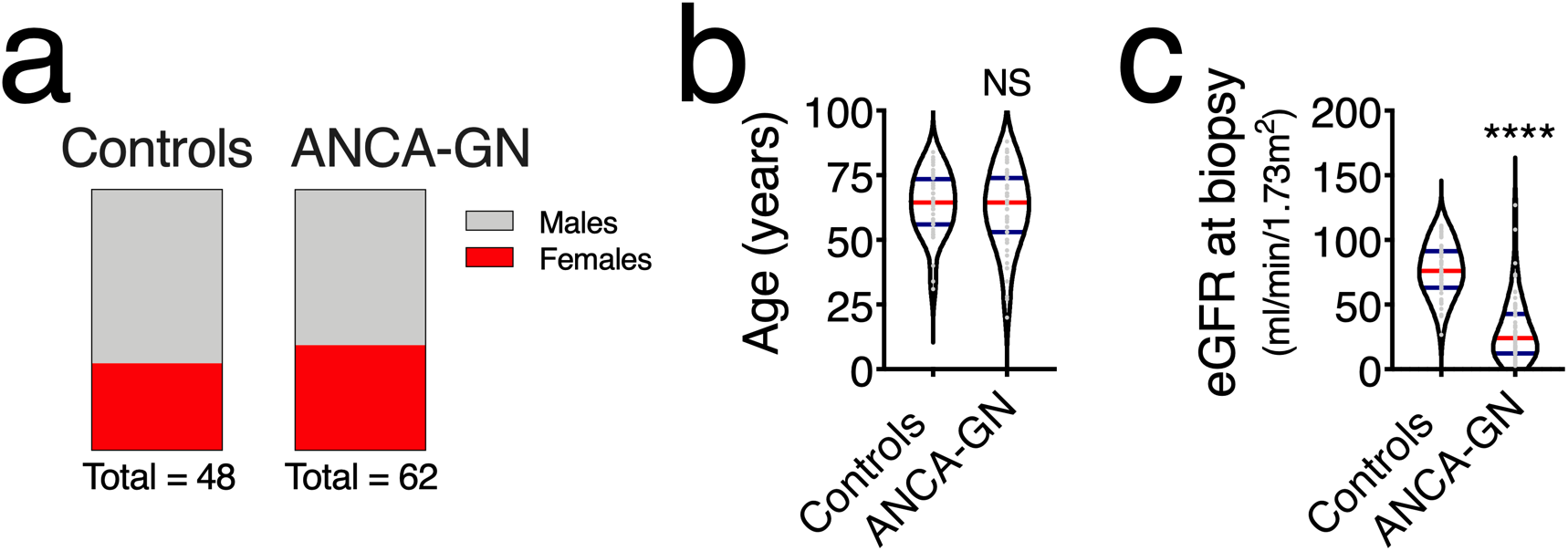
Patient demographics. (a) Similar rates of males and females between controls and ANCA-GN patients. (b) No age differences between controls and ANCA-GN patients. (c) Significant differences in estimated glomerular filtration rate (eGFR) between controls and ANCA-GN patients. In violin plots, each grey dot represents one image, red lines represent medians and blue lines interquartile ranges. ****P<0.0001 and NS: not statistically significant.

**Figure S2.**
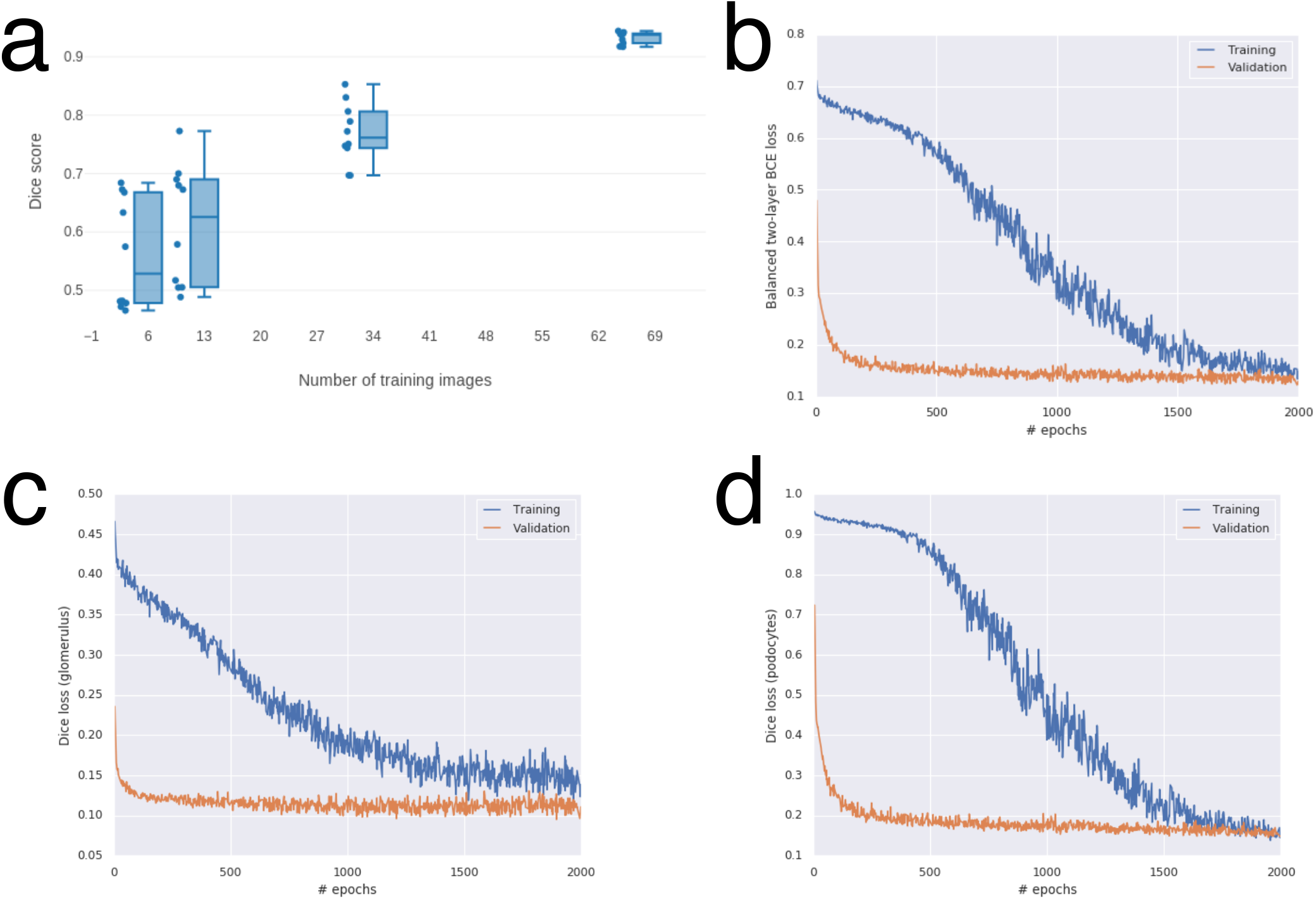
Training of segmentation U-Net. (a) In a 10-fold cross validation, the number of training images was varied. In order to achieve Dice scores >0.90 on a dataset, approximately 65 images were required. The training process was optimised using (b) the balanced 2-layer binary cross entropy (BCE) loss, weighting the individual BCE losses for the glomerulus and podocyte segmentation tasks. The performance was monitored using Dice losses for the (c) glomerulus and (d) podocyte segmentation.

**Figure S3.**
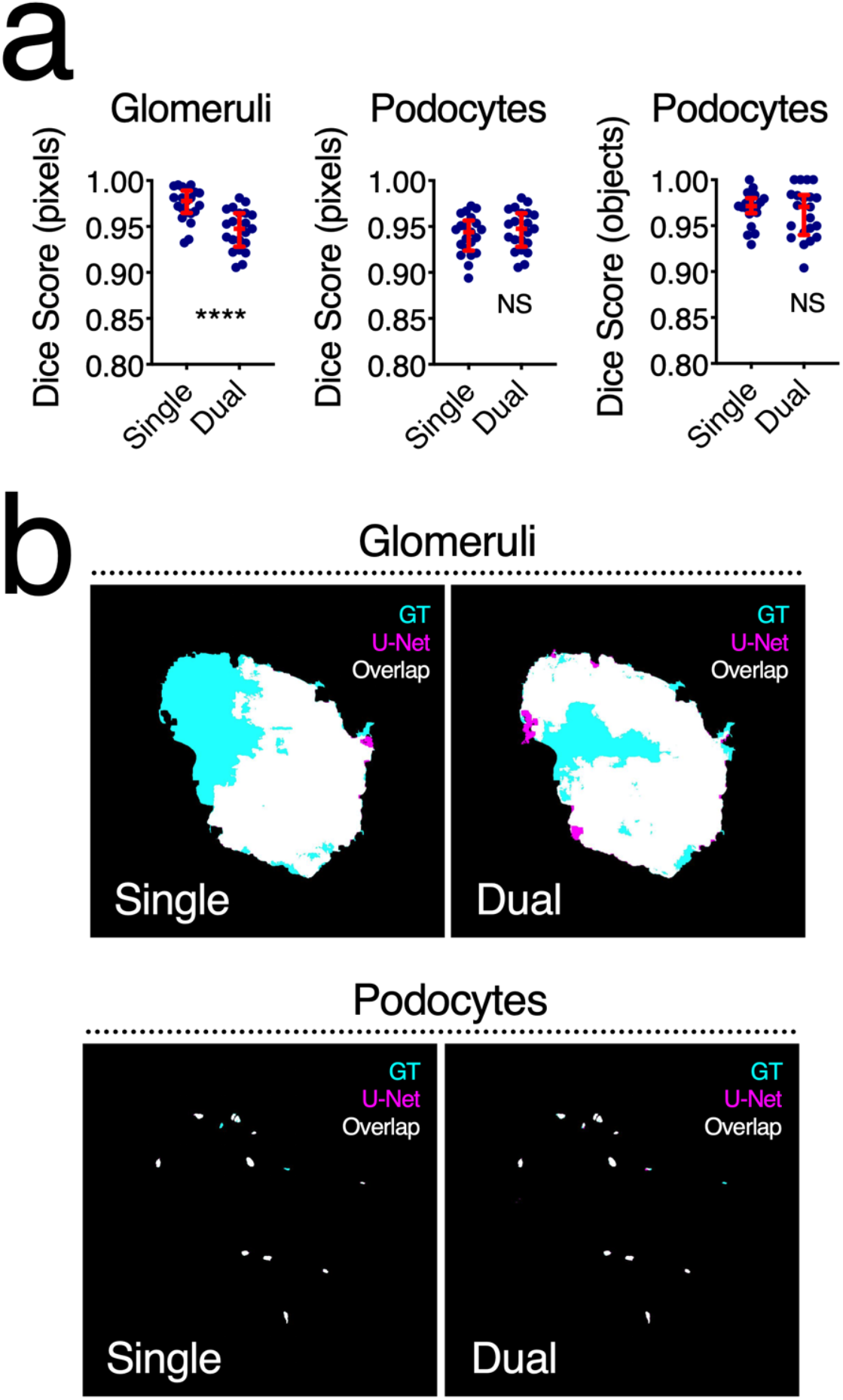
Comparison of single vs. dual output U-Nets. (a) Single and dual segmentation U-Nets provided comparable Dice scores, all over 0.90 for all segmentation tasks. (b) Visual representation of single and dual segmentation U-Nets’ performance. GT: ground truth. Each blue dot represents a single image, red error bars represent medians and interquartile ranges. ****P<0.0001 and NS: not statistically significant.

**Figure S4.**
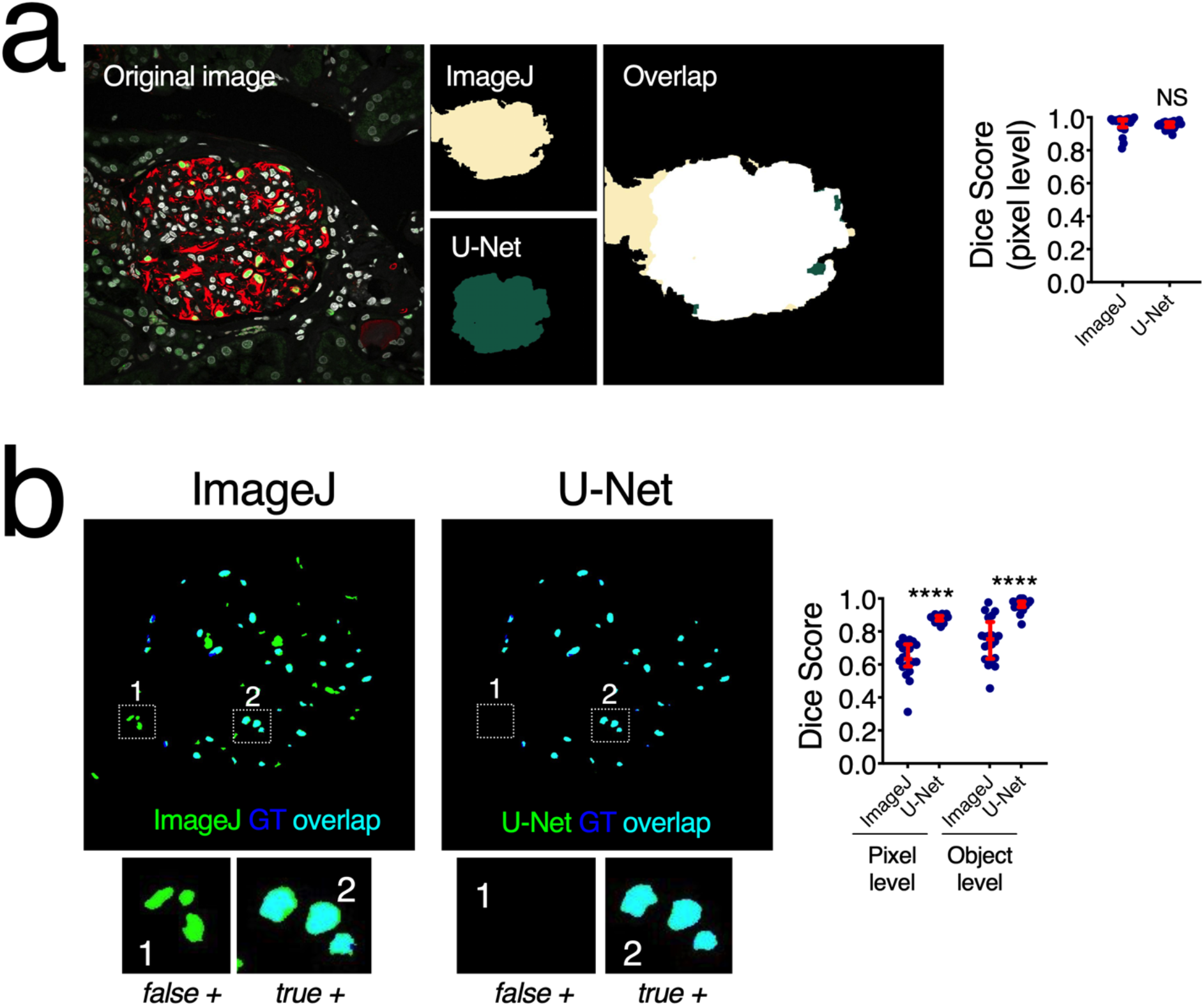
Segmentation U-Net with dual outputs outperforms classical segmentation method. (a) Glomerular segmentation was similar between U-Net and an optimised ImageJ script. However, (b) U-Net significantly outperformed the ImageJ script in podocyte segmentation both at a pixel and object level, reducing the presence of false positives. GT: ground truth. Each blue dot represents a single image, red error bars represent medians and interquartile ranges. ****P<0.0001 and NS: not statistically significant.

**Figure S5.**
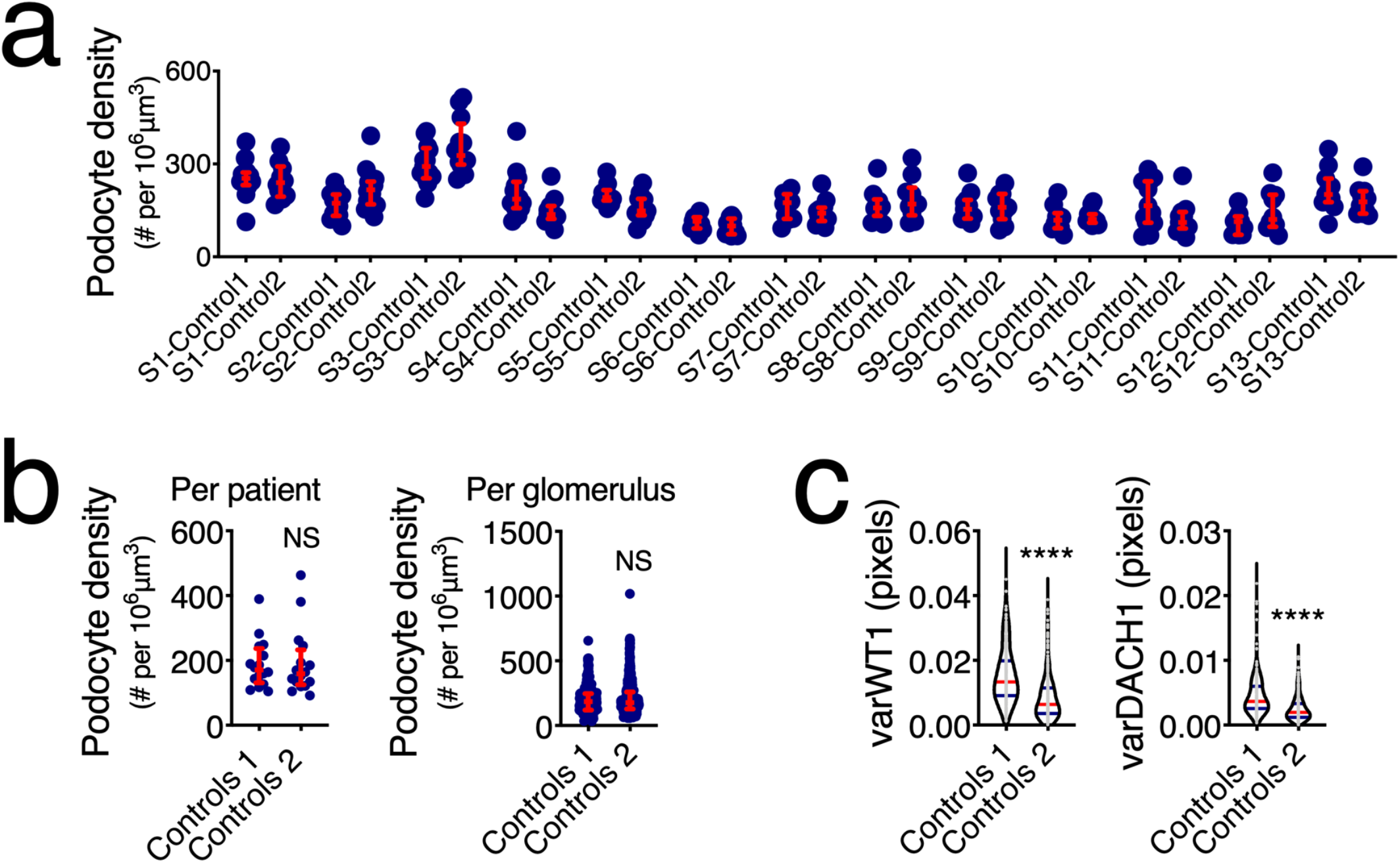
Potential batch effects in external datasets. (a) Direct comparisons of podocyte density within the same subjects. Images were acquired in different sites by different operators using different microscopy systems and consist of 10 randomly sampled glomeruli (out of hundreds available per tissue). None of the comparisons showed statistical significance. (b) Group comparisons using medians per patient or every available glomerulus also showed no statistical differences. (c) Pixel-based analysis did show significant differences for variances in Wilms’ Tumor 1 (WT1) and Dachshund Family Transcription Factor 1 (DACH1). In dot plots, every blue dot represents one glomerulus in (a) and (b-right) and one patient in (b-left), and red error bars represent medians and interquartile ranges. In violin plots, each grey dot represents one image, red lines represent medians and blue lines interquartile ranges. ****P<0.0001 and NS: not statistically significant.

**Figure S6.**
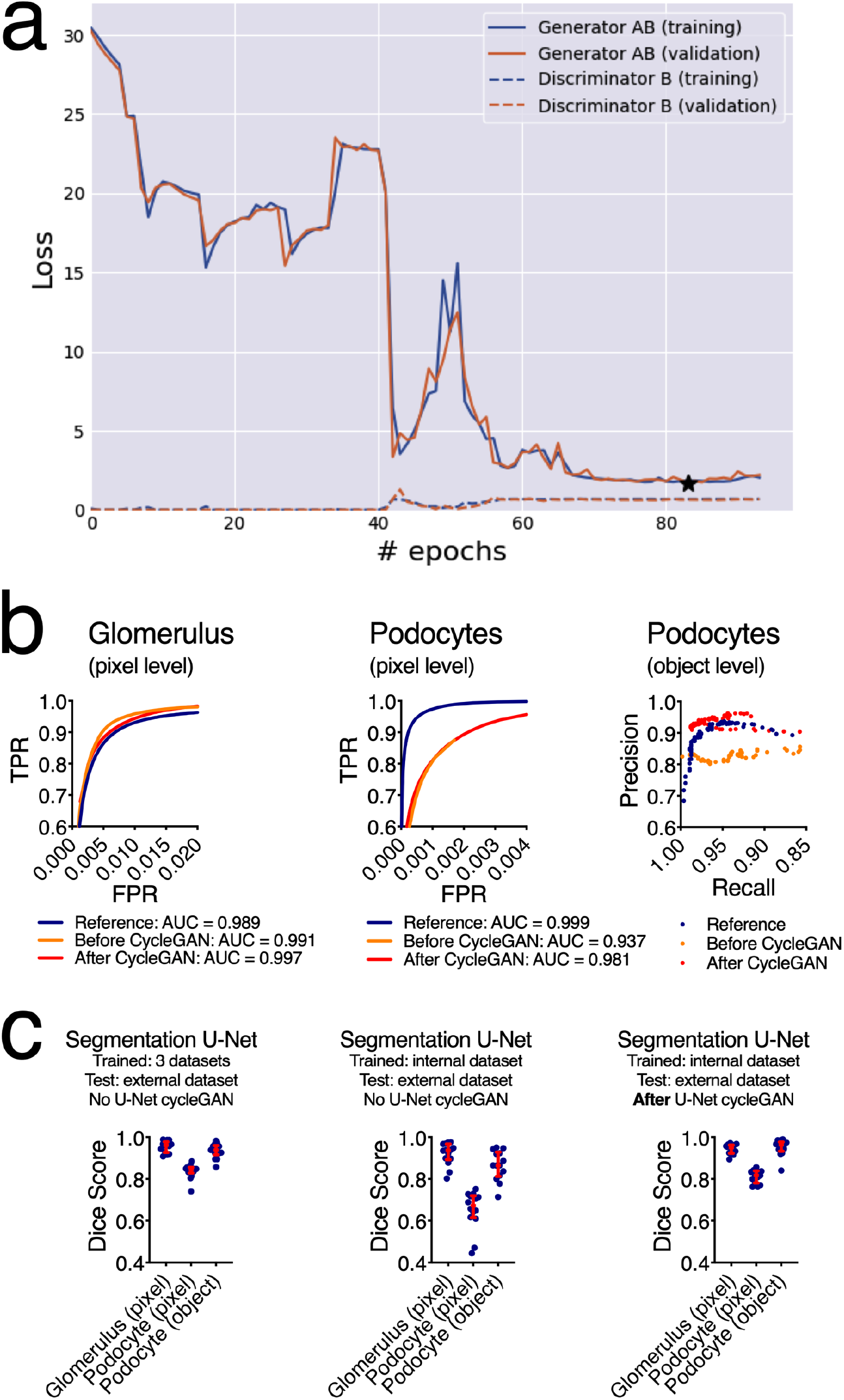
U-Net cycleGAN provides significant improvements in podocyte identification at both pixel and object-level. (a) Training curves for U-Net cycleGAN (the star indicates the best validation loss of the generator, which is the model chosen for evaluation), (b) ROC and precision-recall curves show the performance of segmentation before and after cycleGAN and in relation to the reference dataset. (c) The best results are achieved with manual annotations and re-training of the segmentation U-Net. Using a U-Net cycleGAN leads to comparable Dice scores without the need to re-train the segmentation U-Net. In dot plots, every blue dot represents one image and red error bars represent medians and interquartile ranges.

**Figure S7.**
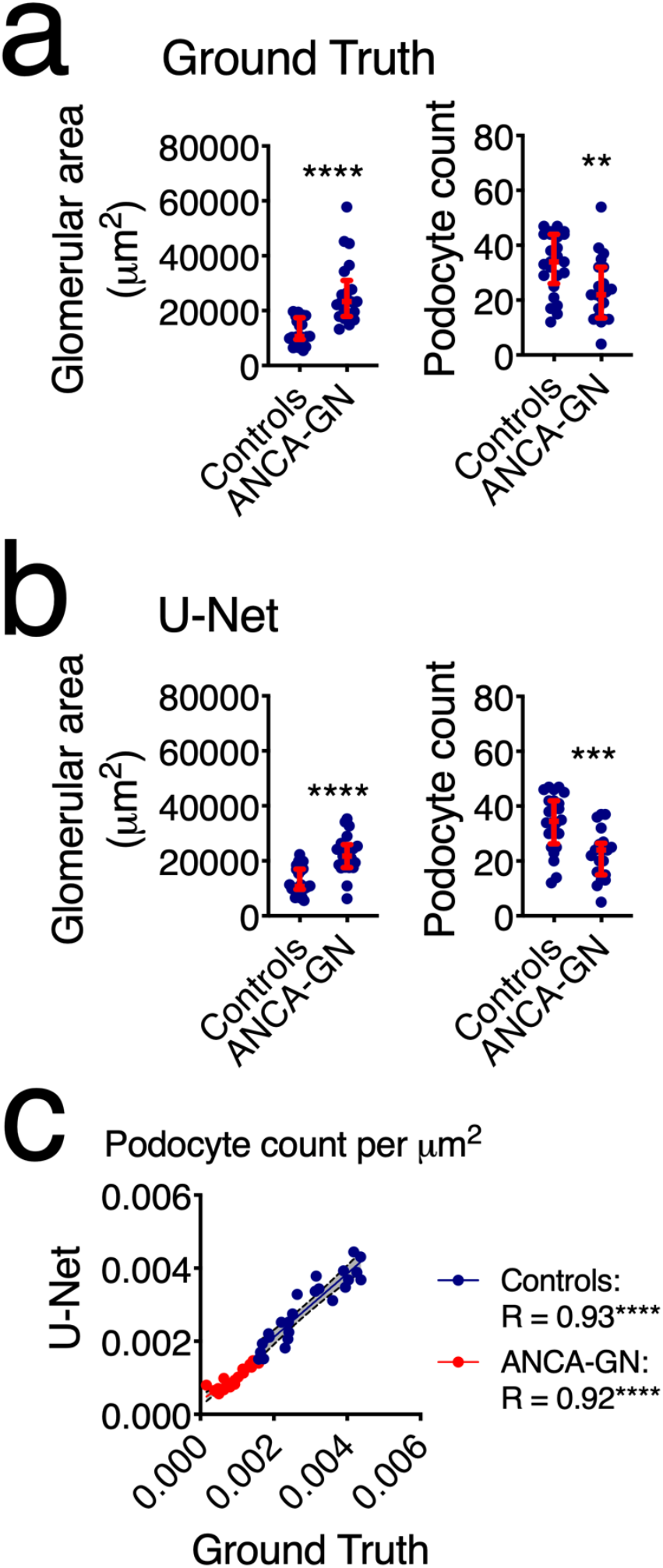
Biological validation of quantitative outputs from segmentation U-Net. (a) 2D quantitative data from manual segmentation (ground truth), including glomerular area, and podocyte count, showing morphometric changes in ANCA-GN patients. (b) 2D quantitative data from segmentation U-Net, including glomerular area and podocyte count, showing identical results to ground truth. (c) Correlation analysis of podocyte count corrected for glomerular area, confirming strong agreement between ground truth and segmentation U-Net for both controls and ANCA-GN datasets. In dot plots, every blue dot represents one image and red error bars represent medians and interquartile ranges. ****P<0.0001, ***P<0.001, and **P<0.01.

**Figure S8.**
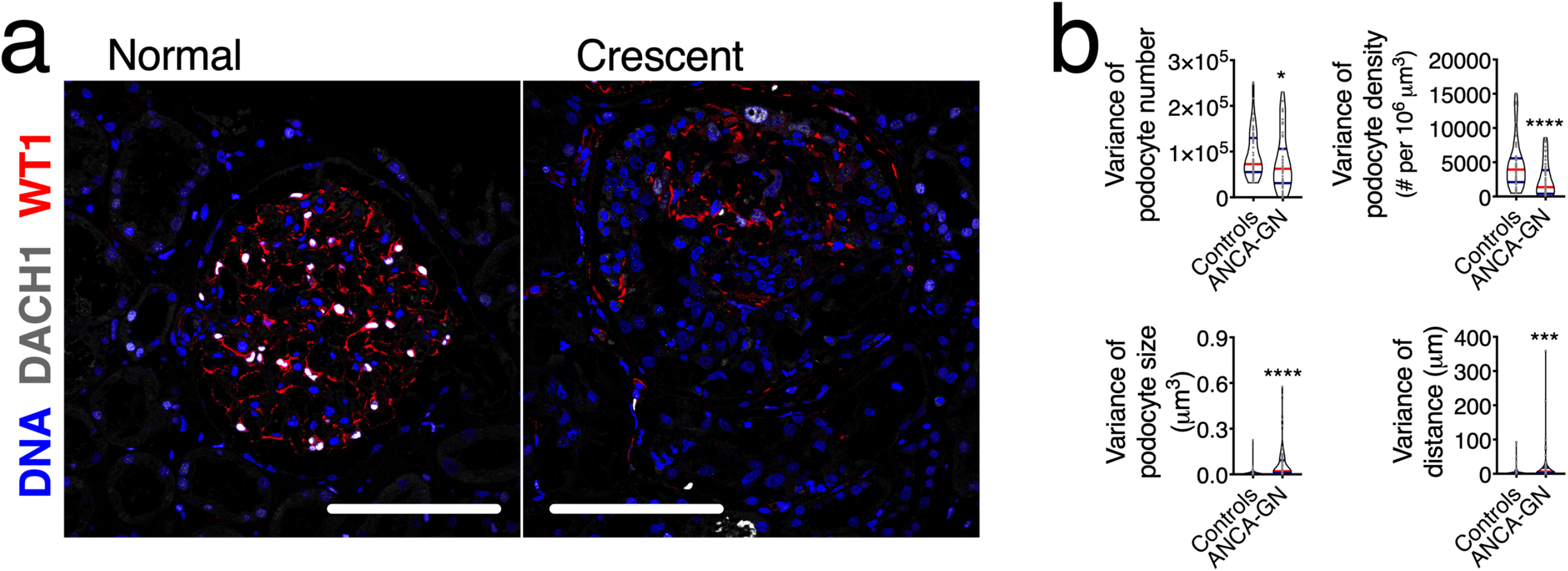
Variation within each biopsy. (a) Representative images of a normal glomerulus (left) and a pathological lesion (crescent, right). (b) Variance of podocyte number, density, size and distances to closest neighbours per subject, highlighting great variability within subjects that is directly affected by the development of kidney disease. Scale bars represent 150μm. ****P<0.0001, ***P<0.001, and *P<0.05.

**Figure S9.**
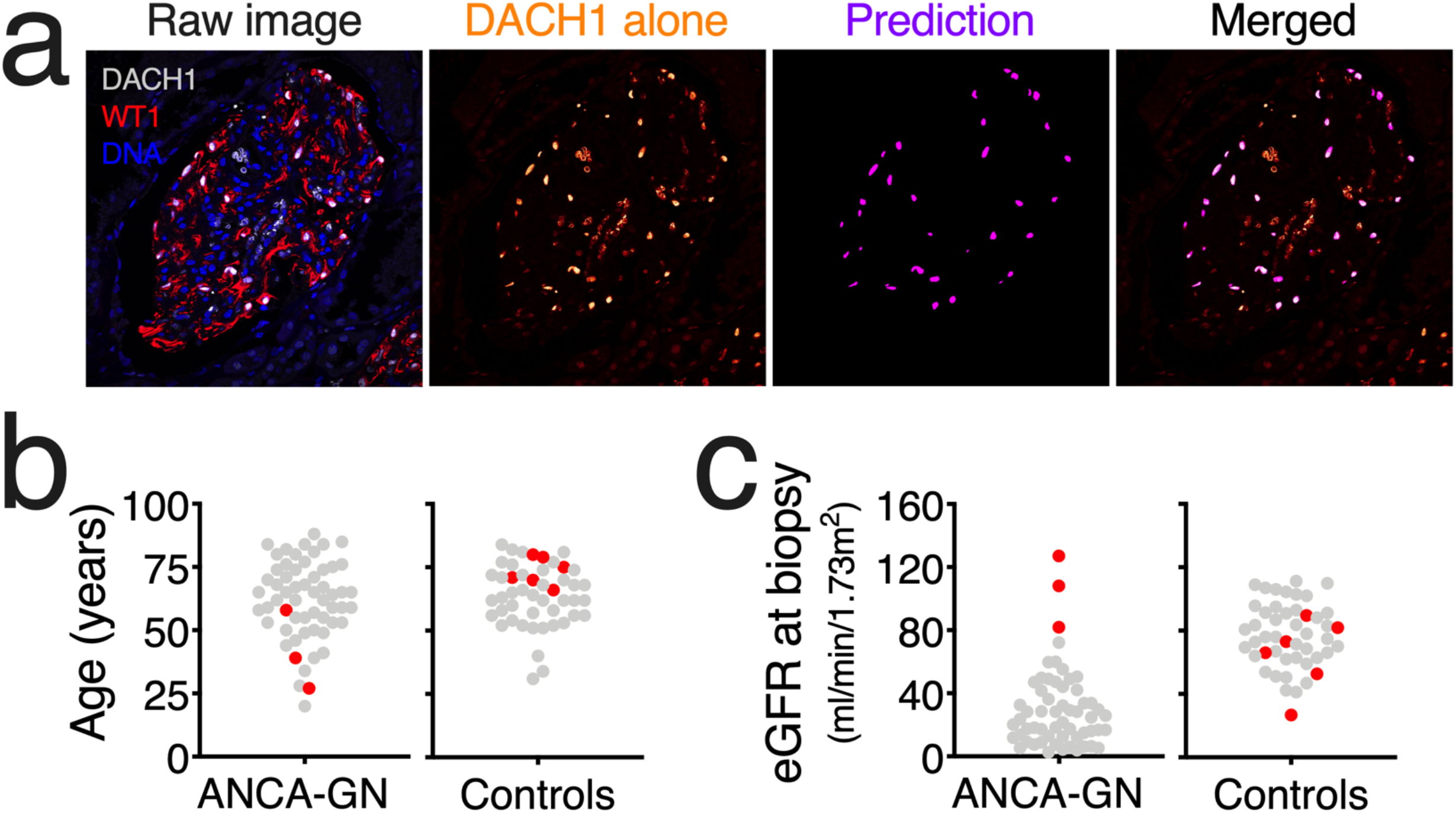
Potential causes of subject misclassification. (a) Representative images of glomerulus from a misclassified subject, showing high prediction accuracy despite of unspecific signal of Dachshund Family Transcription Factor 1 (DACH1). (b) Identification of misclassified subjects (red) in the context of age distributions for ANCA-GN and controls. (c) Identification of misclassified subjects (red) in the context of estimated glomerular filtration rate (eGFR) distributions for ANCA-GN and controls.

